# Impaired cholesterol transport from aged astrocytes to neurons can be rescued by cannabinoids

**DOI:** 10.1101/2023.07.24.550299

**Authors:** Leandro G. Allende, Lautaro Natalí, Andrea B. Cragnolini, Melina M. Musri, Diego de Mendoza, Mauricio G. Martín

## Abstract

Cholesterol is crucial for the proper functioning of eukaryotic cells, especially neurons, which rely on cholesterol to maintain their complex structure and facilitate synaptic transmission. However, brain cells are isolated from peripheral cholesterol by the blood-brain barrier and mature neurons primarily uptake the cholesterol synthesized by astrocytes for proper function. This study aimed to investigate the effect of aging on cholesterol trafficking in astrocytes and its delivery to neurons. Using in vitro and in vivo models of aging, we found that aged astrocytes accumulated high levels of cholesterol in the lysosomal compartment, and this cholesterol buildup can be attributed to the simultaneous occurrence of two events: decreased levels of the ABCA1 transporter which impairs ApoE-cholesterol export from astrocytes, and reduced expression of NPC1, which hinders cholesterol release from lysosomes. We show that these two events are accompanied by increased microR33 in aged astrocytes, which is known to downregulate ABCA1 and NPC1. In addition, we demonstrate that the microR33 increase is triggered by oxidative stress, one of the hallmarks of aging.

By co-culture experiments we also show that aging in vitro impairs the cholesterol delivery from astrocytes to neurons. Remarkably, we found that this altered transport of cholesterol could be alleviated through treatment with endocannabinoids as well as cannabidiol or CBD. Given that reduced neuronal cholesterol affects synaptic plasticity, the ability of cannabinoids to restore cholesterol transport from aged astrocytes to neurons holds significant implications in the field of aging.

## Introduction

Cholesterol is essential for the function of most eukaryotic cells. As a component of the plasma membrane, it plays a major role in determining characteristics such as fluidity, permeability and protein-protein interaction ^[1–3]^. Additionally, as one of the major components of lipid rafts, it also plays an important role in signaling processes and intracellular trafficking ^[4,5]^. Specifically, cholesterol is a key regulator of neuronal function, and any imbalance in cellular cholesterol levels can lead to significant alterations in brain function ^[6–9]^.

The brain contains over 20% of the body’s cholesterol with a substantial portion present in the myelin sheath that insulates axons. Neurons also contain large amounts of cholesterol to maintain their complex morphology and facilitate synaptic transmission. It is important to note that brain cells are isolated from peripheral cholesterol due to the blood-brain barrier, which prevents the entry of cholesterol-rich lipoproteins^[10]^. Within the central nervous system, cholesterol is primarily synthesized by astrocytes and oligodendrocytes. Astrocytes export cholesterol synthesized through ATP binding cassette (ABC) cholesterol transporters, specifically ABCA1, as APOE-cholesterol complexes. Neurons, on the other hand, uptake APOE-cholesterol particles via Lipoprotein Related Receptor 1 (LRP1) mediated endocytosis ^[11]^. The demand for astrocyte-delivered cholesterol appears to be higher in mature neurons compared to young developing cells, as young neurons still synthesize cholesterol to meet the high lipid requirement for membrane expansion and synaptogenesis ^[11]^.

Previous studies from our laboratory have demonstrated that aging of hippocampal neurons is associated with a decrease in membrane cholesterol, leading to reduced synaptic function and memory loss. This occurs through two fundamental mechanisms: alterations in the diffusion and endocytosis of membrane receptors ^[9]^, and the formation of a repressive epigenetic structure in memory gene promoters resulting from impaired neuronal receptor function ^[12]^. Specifically, we have shown that the loss of plasma membrane cholesterol in hippocampal neurons is a consequence of increased expression of the enzyme Cholesterol 24-hydroxylase (CYP46) with aging _[5,13]._

On the other hand, our work with cortical neurons has revealed that aging of these cells leads to lysosomal accumulation of cholesterol due to a decline in the levels of the NPC1 protein, which is involved in the release of cholesterol from neuronal lysosomes following endocytosis ^[14]^. These findings suggest that altered cholesterol redistribution in aged neurons may contribute to suboptimal levels of this lipid in the neuronal plasma membrane.

Considering that mature neurons primarily rely on cholesterol synthesized by astrocytes, this study aimed to investigate whether aging affects the transport of cholesterol from hippocampal astrocytes to hippocampal neurons. This process represents a critical factor in the loss of neuronal cholesterol and may have significant implications for the function of aging neurons.

## Results

### Cholesterol accumulates in the lysosomal compartment in aged astrocytes

Considering a previous study that reported altered cholesterol trafficking in neuronal cells ^[14]^, we first examined whether aging also affects cholesterol trafficking in hippocampal astrocytes. To investigate this, we established an in vitro model of accelerated aging using primary rat astrocytes.

It is well-known that cells grown in a medium containing D-Galactose exhibit an aging phenotype due to the accumulation of oxidative stress^[15]^. Therefore, rat primary hippocampal astrocyte cultures were grown in medium containing 55 mM D-Galactose at different days in vitro to induce accelerated aging. Senescence was assessed by staining for the senescence marker Senescence-Associated β-Galactosidase (SA-β-Gal) ^[16]^. As shown in Figure 1A, astrocytes grown for 5 days in D-Galactose-containing medium exhibited a high proportion of senescent cells positive for SA-β-Gal compared to controls, where no SA-β-Gal staining was observed. Astrocytes grown for 5 days in D-Galactose showed similar proliferative capability and viability to controls (Figures 1B and 1C).

**Figure 1.**
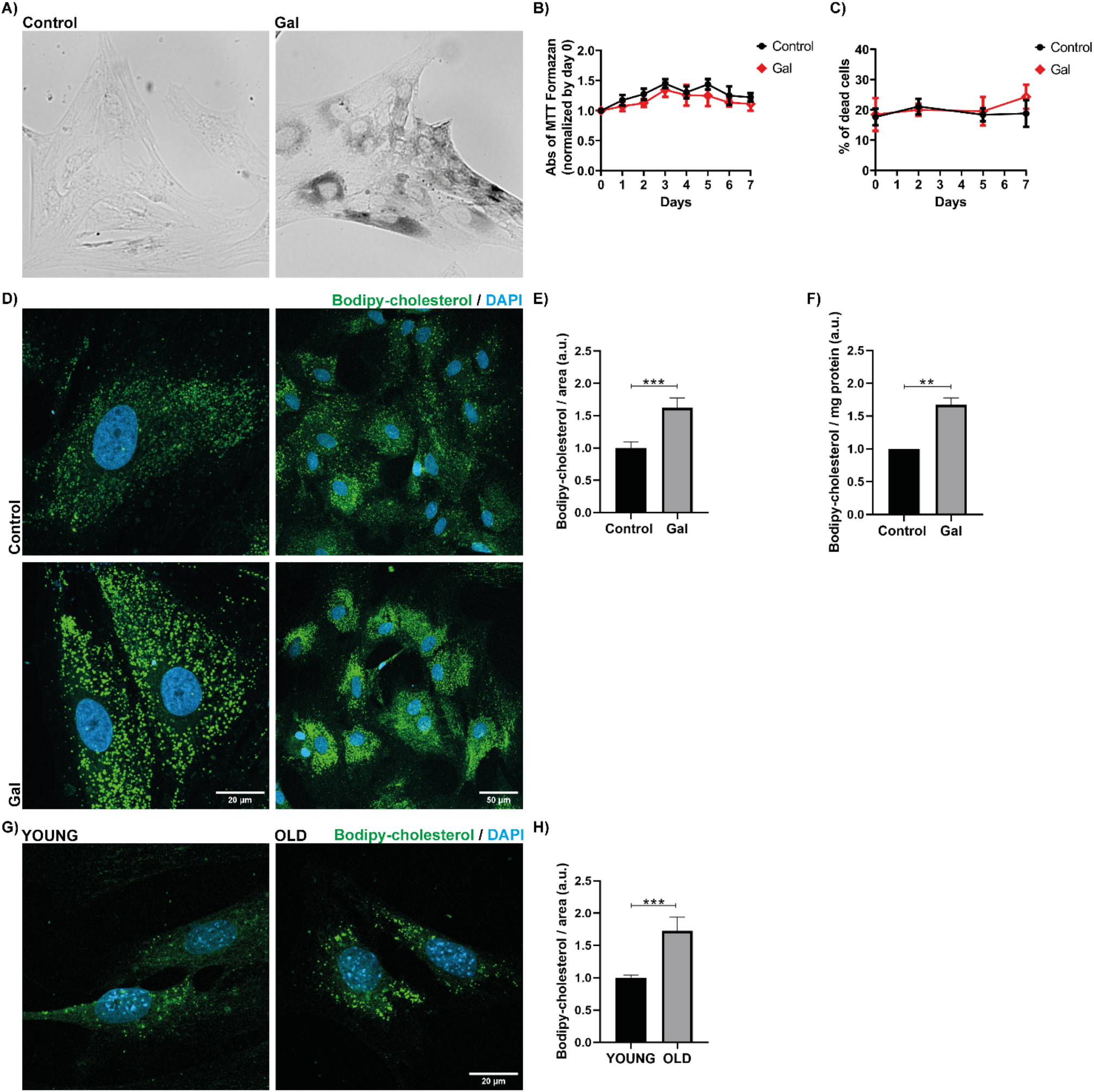
Bodipy-cholesterol accumulation in aged astrocytes. A) Rat hippocampal astrocytes treated with 55mM D-Galactose for 5 days exhibit positive staining for Senescent Associated (SA)-β-galactosidase compared to control cultures. The presence of a dark cytosolic precipitate indicates SA-β-Galactosidase activity. B) MTT assay performed on primary hippocampal astrocytes treated with D-Galactose (GAL) for 7 days shows that cell viability remains unaffected. C) LIVE/DEAD® Viability/Cytotoxicity Assay performed on primary hippocampal astrocytes control or treated with D-Galactose for 7 days reveals a similar percentage of dead cells in both cultures. D) Fluorescence microscopy images demonstrate the accumulation of Bodipy-cholesterol in rat hippocampal astrocytes aged by treatment with D-Galactose. E) Quantification of Bodipy-cholesterol intensity indicates that over 50% of Bodipy-cholesterol accumulates in aged astrocytes in vitro compared to controls. F) Fluorometric measurements of Bodipy-cholesterol in total extracts of control or D-Galactose treated astrocytes exhibit significant cholesterol accumulation in aged cells. G) Fluorescence microscopy images reveal the accumulation of Bodipy-cholesterol in hippocampal astrocytes obtained from 20-month-old mice (OLD) compared to astrocytes from 2-month-old mice (YOUNG). H) Quantification of the images shown in G indicates that over 50% of Bodipy-cholesterol accumulates in primary astrocytes prepared from old mice. Data are presented as Mean ± SEM. ***P-value < 0.001, **P-value < 0.01.

To examine the distribution of cholesterol in control and aged astrocytes, the cultures were incubated overnight with the fluorescent cholesterol analog Bodipy-cholesterol and analyzed using fluorescence confocal microscopy (Figure 1D). The results in Figures 1D and 1E demonstrate that aged astrocytes accumulated over 50% higher levels of cholesterol compared to controls (Control = 1.0000 ± 0.0916 SEM, aged = 1.6217 ± 0.1516 SEM, p = 0.0009). This finding was further confirmed by fluorometric analysis of total extracts from control and aged rat primary astrocytes, which showed a 67% increase in cholesterol accumulation in aged astrocytes relative to controls (Figure 1F, Control = 1.0000, aged = 1.6700 ± 0.1059 SEM). Cholesterol accumulation was also observed in aged human primary astrocytes in vitro (Supp. Fig. 1A and 1B).

To validate our model of accelerated aging, we assessed Bodipy-cholesterol accumulation in primary astrocytes prepared from the hippocampus of young (2 months) or old (20 months) mice. Once again, we observed over 50% cholesterol accumulation in old astrocytes compared to young ones (Young = 1.0000 ± 0.0418 SEM, Old = 1.7231 ± 0.2144 SEM, p = 0.0008; Figures 1G and 1H), indicating a cholesterol trafficking defect in aged astrocytes.

Next, we determined the intracellular compartment where cholesterol accumulates in aged astrocytes through colocalization studies between Bodipy-cholesterol and markers such as Lamp2 (lysosomal marker), Rab5 (early endosome marker), or Rab11 (late endosome marker). Control or aged primary astrocytes were loaded with Bodipy-cholesterol overnight and immunolabeled for each of these markers. Analysis of immunofluorescence confocal images revealed a significant increase in Bodipy-cholesterol-Lamp2 colocalization in aged astrocytes (Figure 2A and Figure 2B), with no changes observed in Bodipy-cholesterol-Rab5 or Rab11 colocalization (Supp. Figure 2A and Supp. Figure 2B).

**Figure 2.**
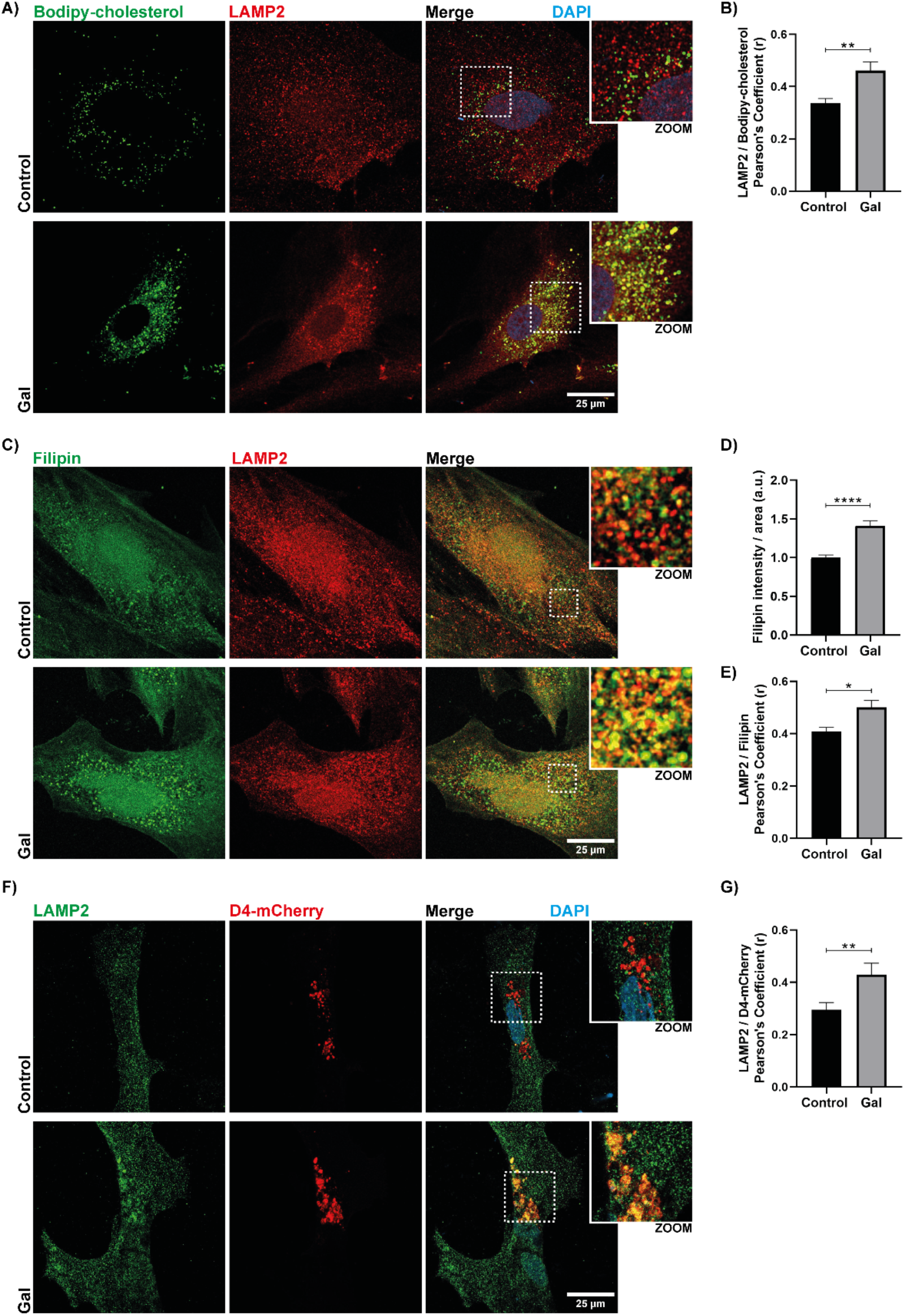
Cholesterol accumulation in the lysosomal compartment in astrocytes aged in vitro. A) Immunofluorescence confocal images of control or D-Galactose-treated rat hippocampal astrocytes (GAL) labeled with Bodipy-cholesterol and the lysosomal marker LAMP2. B) Quantitative analysis of images including those shown in A, reveals increased colocalization between Bodipy-cholesterol and LAMP2 in aged astrocytes. LAMP2 / Bodipy-cholesterol Pearson’s Coefficient (r): Control = 0.3364 ± 0.0185SEM, Gal = 0.4619 ± 0.0328SEM (p=0.0079). C) Immunofluorescence images of rat hippocampal astrocytes under control or aged conditions, labeled with Filipin to visualize endogenous cholesterol and the lysosomal marker LAMP2. D) Quantification of Filipin intensity in control or D-Galactose-treated astrocytes demonstrates the accumulation of endogenous cholesterol in aged astrocytes. E) Analysis of images including those shown in C indicates increased colocalization between Filipin and LAMP2 in aged astrocytes. LAMP2 / Filipin Pearson’s Coefficient (r): Control = 0.4089 ± 0.0152SEM, Gal = 0.5006 ± 0.0271SEM (p=0.0242). F) Immunofluorescence images of rat hippocampal astrocytes under control or aged conditions, expressing the D4-mCherry probe to visualize endogenous cholesterol and the lysosomal marker LAMP2. G) Analysis of images including those shown in F reveals increased colocalization between D4-mCherry and LAMP2 in aged astrocytes. LAMP2 / D4-mCherry Pearson’s Coefficient (r): Control = 0.2956 ± 0.0271SEM, Gal = 0.4300 ± 0.0433SEM (p=0.0093). Data are presented as Mean ± SEM. ****P-value < 0.0001, ***P-value < 0.001, **P-value< 0.01, *P-value < 0.05.

Bodipy-cholesterol is a probe that, once added to astrocytes, equilibrates within the cell with endogenous cholesterol. Therefore, we hypothesized that endogenous cholesterol accumulates in the lysosomes of aged astrocytes. To confirm these results, we examined endogenous cholesterol distribution using two other methods: filipin staining and D4-mCherry labeling. Filipin is a fluorescent polyene macrolide antibiotic isolated from *S. filipinensis* that binds to cholesterol and is widely used as a probe for determining cholesterol location in biological membranes. D4 is the cholesterol binding domain of perfringolysin O (θ toxin) from *Clostridium perfringens* fused to the red fluorescence protein mCherry ^[17]^.

Consistent with the Bodipy-cholesterol results, filipin staining of control and aged primary astrocytes revealed a 41% cholesterol accumulation in aged cells (Control = 1.0000 ± 0.0345 SEM, aged = 1.4132 ± 0.0635 SEM, p < 0.0001; Figures 2C and 2D) that colocalized with LAMP2-positive compartments (Figures 2C and 2E). Similar results were obtained when visualizing cholesterol distribution in astrocytes expressing the D4-mCherry fusion protein. As shown in Figure 2G, this probe accumulated in lysosomal compartments in aged astrocytes, as indicated by the increased colocalization of LAMP2 and D4-mCherry.

Taken together, our results indicate that cholesterol accumulates in the lysosomal compartment of aged astrocytes.

### Cholesterol accumulation in aged astrocytes is a consequence of downregulation of NPC1 and ABCA1

The release of cholesterol from the endolysosomal compartment is mediated by the cooperative action of two proteins, Niemann-Pick Type C Protein 1 and 2 (NPC1 and NPC2), which facilitate the incorporation of cholesterol into the intracellular pool. Alterations in both NPC1 and NPC2 are implicated in the neurodegenerative lysosomal disorder known as Niemann-Pick Type C (NPC) disease ^[18]^, characterized by the accumulation of non-esterified cholesterol and sphingolipids in late endosomes/lysosomes ^[19]^. Recent research by Guix et al. ^[14]^ demonstrated that the NPC1 gene is downregulated in cortical neurons aged in vitro due to increased expression of microRNA 33 (miR-33). miR-33 controls cholesterol homeostasis under conditions of low cholesterol by reducing the expression of the ABCA1 and ABCG1 genes, as well as the NPC1 gene, to minimize cholesterol efflux ^[20]^.

We investigated the possibility that cholesterol accumulation in aged astrocytes could be attributed to NPC1 downregulation. According to this hypothesis, decreased levels of NPC1 protein (0.7368 ± 0.0779 SEM, p = 0.0149) and mRNA (0.6732 ± 0.0791 SEM, p = 0.0258) were observed in aged primary hippocampal astrocytes compared to controls (Figure 3A and 3B). Consistent with these findings, reduced NPC1 protein and mRNA levels were also detected in the hippocampal tissue of 20-month-old mice compared to 2-month-old individuals (Figure 3C and 3D). Thus, our results demonstrate that aging triggers a Niemann-Pick phenotype not only in astrocytes aged in vitro but also in vivo in the hippocampus of old mice.

**Figure 3.**
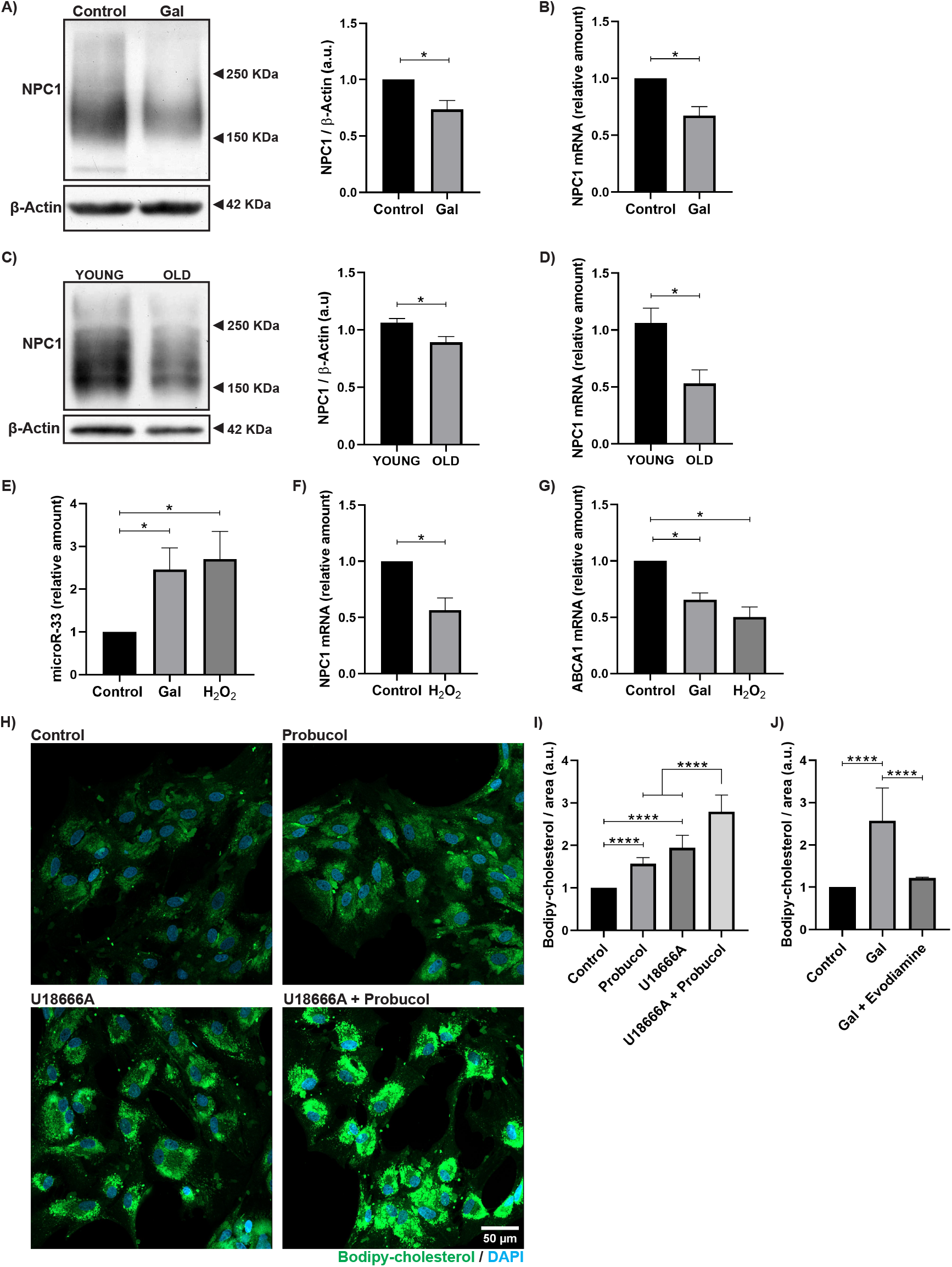
NPC1 and ABCA1 downregulation leads to remarkable cholesterol accumulation in aged astrocytes. A) Western blot analysis, along with quantification showing decreased NPC1 protein levels in aged primary hippocampal astrocytes compared to control cultures. B) RT-qPCR analysis showing decreased NPC1 mRNA levels aged rat primary hippocampal astrocytes. C) Western blot analysis, along with quantification showing decreased NPC1 protein levels in the hippocampus of old (20 months-old) mice compared to young (2 months-old) C57BL/6 mice (YOUNG = 1.0630 + 0.0364 SEM, OLD = 0.8919 + 0.0494 SEM, p = 0.0303). D) RT-qPCR analysis showing decreased NPC1 mRNA levels in the hippocampus of old C57BL/6 mice compared to young individuals (YOUNG = 1.0635 + 0.1290 SEM, OLD = 0.5290 + 0.1213 SEM, p = 0.0242). E) Quantification of miR-33 levels by RT-qPCR in control, aged, or H2O_2_-treated hippocampal astrocytes, indicating increased expression of miR-33 in these cells due to aging or oxidative stress. F) RT-qPCR analysis showing decreased NPC1 mRNA levels in hippocampal astrocytes treated with H_2_O_2_ compared to untreated controls. G) RT-qPCR quantification of ABCA1 mRNA levels in control, aged, or H_2_O_2_-treated hippocampal astrocytes, demonstrating downregulation of ABCA1 in primary astrocytes due to aging or oxidative stress. H) Fluorescence microscopy images of rat primary hippocampal astrocytes treated with Probucol 10µM (ABCA1 inhibitor), U18666A 4µg/mL (NPC1 inhibitor), and the combination of both, labeled with Bodipy-cholesterol. I) Quantification of images including those shown in H, indicating that NPC1 or ABCA1 inhibition leads to Bodipy-cholesterol accumulation in astrocytes, with enhanced effects observed when both proteins are inhibited. J) Treatment of aged astrocytes with the ABCA1 agonist evodiamine leads to a significant decrease in the levels of accumulated Bodipy-cholesterol. Data are represented as Mean ± SEM. ****P-value < 0.0001, *P-value < 0.05.

To analyze if the upregulation of miR-33 underlies the decreased expression of NPC1 in aged astrocytes, the levels of this miRNA were measured in control or D-Galactose-treated primary cultures. The results in Figure 3E clearly show a 2.4550 ± 0.5087 SEM (p = 0.0444) fold increase in miR-33 levels in aged astrocytes compared to controls, providing a mechanistic insight into NPC1 downregulation in this model. Consistent with these results, miR-33 levels were found to be increased in the hippocampus of 2-month-old and 20-month-old mice, indicating increased miR-33 expression during aging in vivo (Supp. Figure 3).

Considering that D-Galactose treatment promotes the accumulation of oxidative stress, which is one of the hallmarks of aging, we tested the possibility that oxidative stress triggers miR-33 expression in astrocytes. As shown in Figure 3E, a significant increase in miR-33 levels of 2.7050 ± 0.6462 SEM (p = 0.0356) fold over controls was observed in astrocytes treated with H_2_O_2_. These findings suggest that, in addition to low cholesterol conditions, the accumulation of oxidative stress may induce NPC1 decay and alter cholesterol trafficking in astrocytes via miR-1. 33. Consistent with these data, the addition of H_2_O_2_ to rat primary astrocytes resulted in a reduction in NPC1 mRNA levels to 0.5640 ± 0.1106 SEM (p = 0.0169) fold over control (Figure 3F).

Once cholesterol is synthesized in astrocytes, it is exported to the extracellular medium as ApoE-cholesterol lipoprotein complexes through ABCA1 transporters ^[11]^. Considering that the ABCA1 gene is also a target of miR-33, we investigated whether mRNA-ABCA1 levels decrease in aged astrocytes. The RT-qPCR results showed significantly reduced levels of mRNA-ABCA1 in aged astrocytes (0.6541 ± 0.0356 SEM, p = 0.0104) compared to controls (Figure 3G). As expected, treatment of primary astrocytes with H_2_O_2_ further decreased ABCA1 gene expression (0.5031 ± 0.0510 SEM, p = 0.0104) compared to controls (Figure 3G).

To confirm that reduced NPC1 and ABCA1 are sufficient conditions to lead to cholesterol accumulation in astrocytes, we used a pharmacological approach to inhibit NPC1, ABCA1, or both. In these experiments, rat primary astrocytes were treated with U18666A to inhibit NPC1 and Probucol to inhibit ABCA1, and the accumulation of Bodipy-cholesterol was measured using fluorescence microscopy. Our results showed that NPC1 inhibition led to a 1.9459 ± 0.2929 SEM (p < 0.0001) cholesterol accumulation compared to controls, while ABCA1 inhibition resulted in a 1.5696 ± 0.1450 SEM (p < 0.0001) cholesterol accumulation (Figures 3H and 3I). As expected, the combined effect of ABCA1 and NPC1 inhibition was observed, as cells treated with U18666A + Probucol accumulated significantly higher cholesterol levels (2.7908 ± 0.3940 SEM, p < 0.0001) compared to astrocytes treated with individual inhibitors or untreated controls (Figures 3H and 3I). Further supporting the notion that impaired cholesterol export through ABCA1 transporters contributes to cholesterol accumulation in aged astrocytes, we found that the high levels of Bodipy-cholesterol in aged astrocytes decreased upon treatment with the ABCA1 agonist Evodiamine (Gal = 2.5710 ± 0.77285 SEM; Gal + Evodiamine = 1.2150 ± 0.0192 SEM, p < 0.0001, Figure 3J).

Taken together, our results indicate that oxidative stress accumulated during aging can trigger a Niemann-Pick phenotype in old astrocytes through the interplay of two simultaneous events: impaired ApoE-cholesterol export due to decreased ABCA1 levels and impaired cholesterol release from lysosomes due to reduced NPC1 expression.

### Aged astrocytes exhibit impaired cholesterol delivery to neurons

As mentioned previously, mature neurons rely heavily on cholesterol synthesized by astrocytes. Our data indicate that cholesterol accumulation occurs in old astrocytes, partly due to decreased expression of ABCA1 transporters, suggesting that cholesterol delivery from astrocytes to neurons is impaired during aging. To investigate this possibility, we examined cholesterol transport from astrocytes to neurons using primary co-cultures of these two cell types. In these experiments, rat primary astrocytes, either untreated or treated with D-Galactose, were loaded overnight with Bodipy-cholesterol. The cells were then washed with 1x Hanks buffer to remove any unincorporated Bodipy-cholesterol, and the medium was replaced with Neurobasal neuronal medium. Glass coverslips containing primary hippocampal neurons were subsequently placed on the dishes containing either control or aged astrocytes loaded with Bodipy-cholesterol (Figure 4A). Direct contact between astrocytes and neurons was disrupted by applying paraffin dots on the glass coverslips. After 6 hours of incubation, the incorporation of Bodipy-cholesterol derived from astrocytes into neuronal cells was measured using fluorescence confocal microscopy.

**Figure 4.**
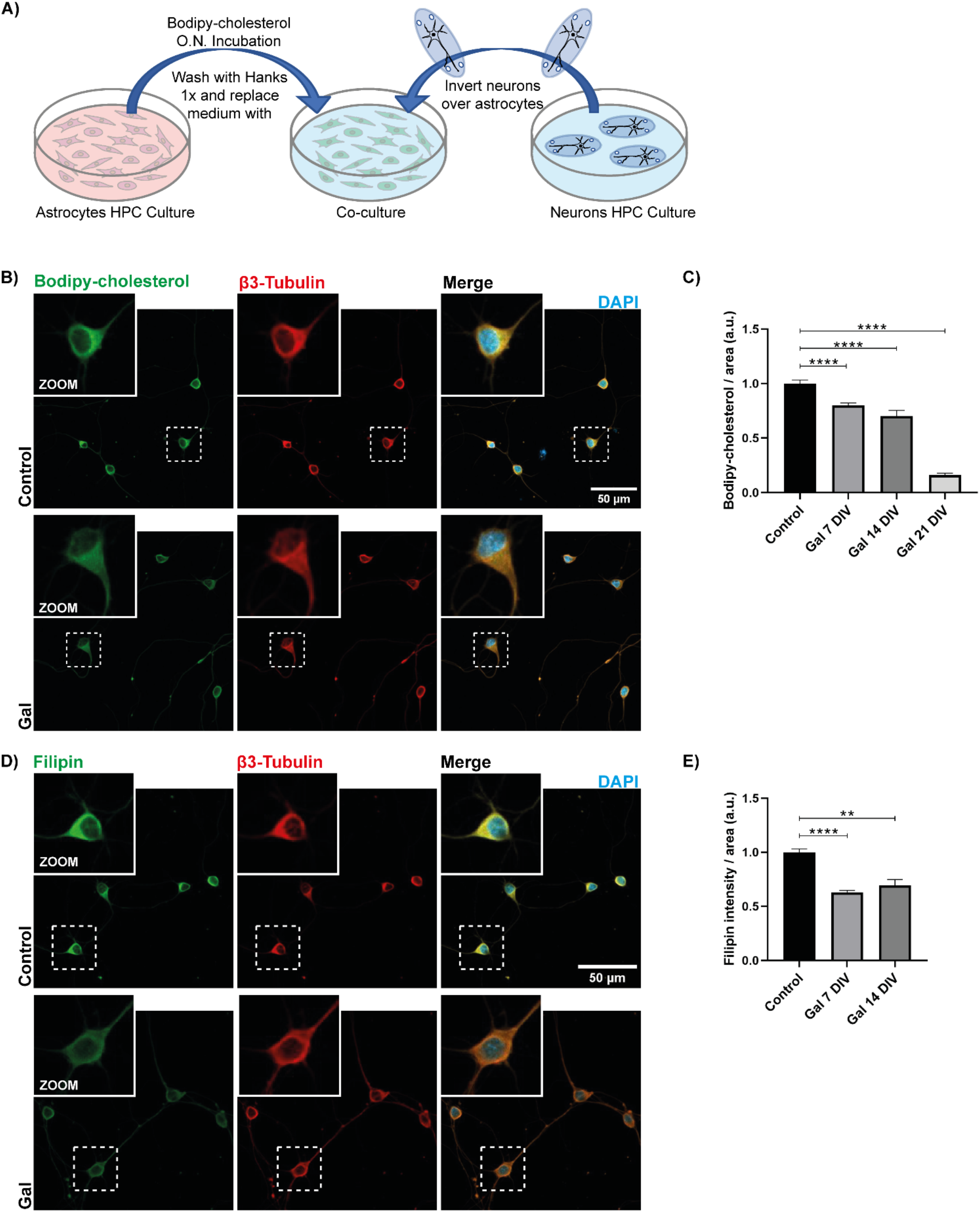
Aging impairs cholesterol transport from astrocytes to neurons. A) Schematic diagram illustrating the co-cultures of rat hippocampal neurons with primary astrocytes. B) Immunofluorescence images showing the uptake of Bodipy-cholesterol by 7DIV hippocampal neurons co-cultured with either control or aged astrocytes for 6 hours. Neuronal cells were labeled with anti-β3-Tubulin antibodies. C) Quantification of Bodipy-cholesterol incorporation in 7DIV, 14DIV, and 21DIV neurons incubated with aged astrocytes compared to young controls. D) Similar to experiment described in B, cholesterol incorporated by 7DIV hippocampal neurons was labeled with Filipin. E) Filipin intensity quantification to determine the cholesterol levels incorporated in 7DIV and 14DIV neurons incubated for 6 hours with aged astrocytes compared to young controls. The plots in C and E illustrate that neurons incubated for 6 hours with aged astrocytes incorporate lower levels of cholesterol compared to neurons incubated with young cultures, and this effect becomes more pronounced as neurons age from 7DIV to 21DIV. (DIV) Days in vitro. Data are represented as Mean ± SEM. ****P-value < 0.0001, **P-value < 0.01.

As depicted in Figures 4B and 4C, lower levels of Bodipy-cholesterol were observed in neurons co-cultured with aged astrocytes compared to neurons incubated with control cultures. As expected, this effect was more pronounced in neurons at 14 and 21 days in vitro (DIV), which highly depend on astrocytic cholesterol, whereas the differences were less prominent (although still significant) in 7DIV neurons (Figure 4C: 7DIV = 0.8023 ± 0.0199 SEM, 14DIV = 0.7013 ± 0.0526 SEM, 21DIV = 0.1623 ± 0.0154, p < 0.0001). Similar results were obtained when the same experiment was repeated by staining the neurons with filipin to measure the uptake of cholesterol endogenously synthesized by control or aged astrocytes (Figures 4D and 4E).

Taken together, our results indicate that cholesterol transport from astrocytes to neurons is impaired during aging.

### Cholesterol accumulation in aged astrocytes is reversed by treatment with cannabinoids

Experiments conducted by Galles et al. ^[21]^ indicated that endocannabinoids can promote cholesterol recycling in a *C. elegans* mutant for the ncr1 and ncr2 genes, which are orthologous to the human Niemann Pick C disease gene npc1. Similar results were obtained by Bartoll et al. ^[22]^, who demonstrated that inhibition of Fatty Acid Hydrolase (FAAH), involved in endocannabinoid breakdown, reduces sphingomyelin and cholesterol accumulation in NPC models.

Considering these findings, we aimed to evaluate whether endocannabinoids can rescue the alterations in cholesterol traffic observed in aged astrocytes. In these experiments, control or aged astrocytes were loaded with Bodipy-cholesterol overnight and then incubated for 6 hours in the presence of the endocannabinoids anandamide (AEA) or 2-acylglycerol (2-AG). As shown in Figures 5A and 5B, endocannabinoids effectively reverse the lysosomal cholesterol accumulation observed in aged astrocytes (Control = 1.0000 ± 0.0238 SEM, GAL = 1.9630 ± 0.0813 SEM, GAL + AEA = 1.2760 ± 0.0418 SEM, GAL + 2-AG = 1.2485 ± 0.0549 SEM, p < 0.0001). Interestingly, the phytocannabinoid CBD also reverses the cholesterol accumulation observed in aged astrocytes, as demonstrated in the plot in Figure 5B (GAL + CBD = 1.1832 ± 0.0577 SEM, p < 0.0001).

**Figure 5.**
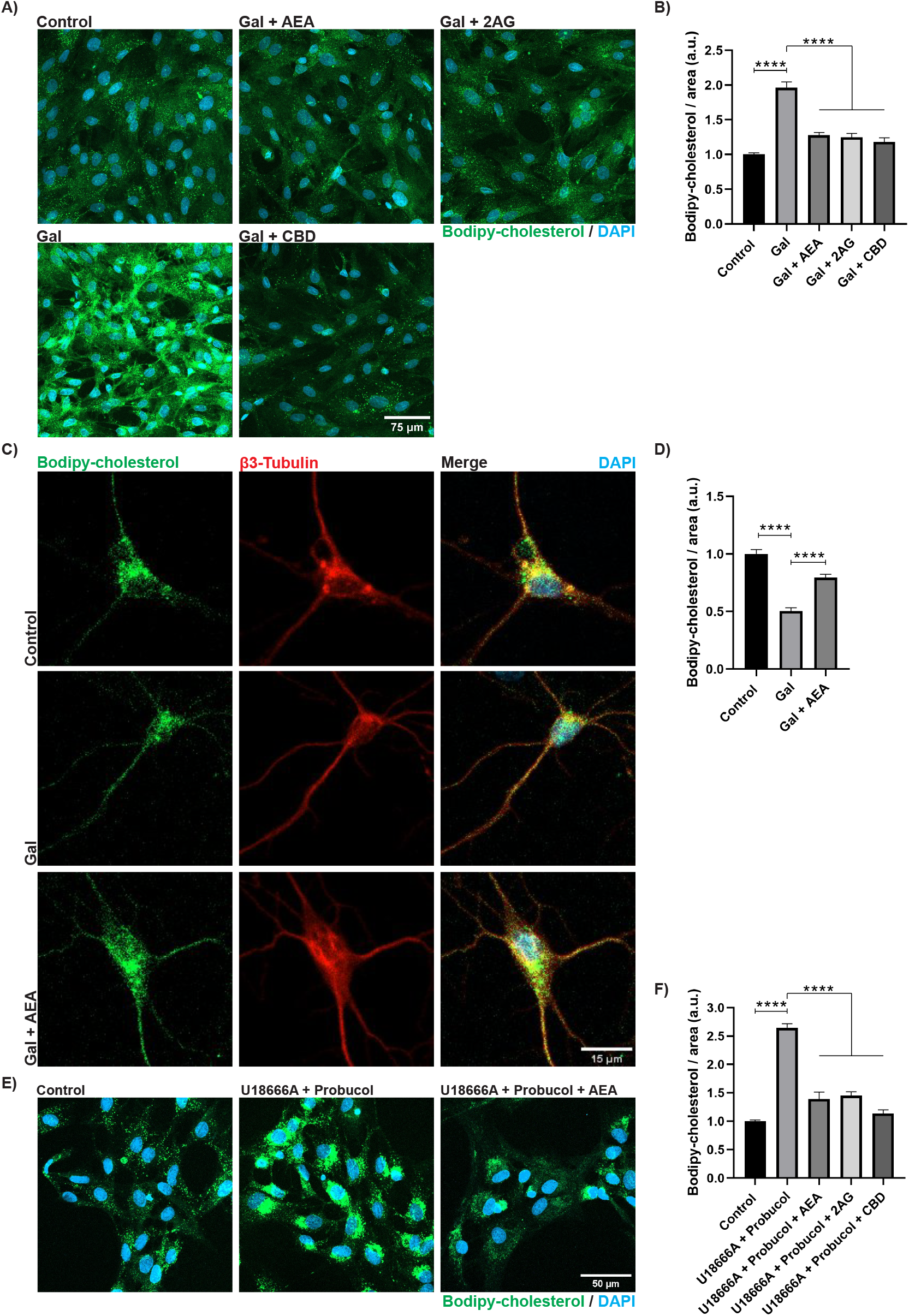
Cannabinoids rescue the accumulation of cholesterol in aged astrocytes and its transport to neurons. A) Fluorescence microscopy images of aged rat hippocampal astrocytes labeled with Bodipy-cholesterol and treated with the cannabinoids Anandamide (AEA 2µM), 2-Arachidonoylglycerol (2AG 2µM) and cannabidiol (CBD 5µM). B) Quantification of Bodipy-cholesterol intensity from images as shown in A demonstrates that treatment of aged astrocytes with AEA, 2AG, or CBD significantly reduces the Bodipy-cholesterol accumulation characteristic of aged astrocytes. C) Immunofluorescence images showing the levels of Bodipy-cholesterol incorporated in 7DIV hippocampal neurons co-cultured with either control, aged, or aged astrocytes in the presence of AEA for 6 hours. Neuronal cells were labeled for β3-Tubulin. D) Quantification of Bodipy-cholesterol intensity from images including those in C demonstrates that the cholesterol uptake by neurons co-cultured with aged astrocytes is improved by addition of AEA. E) Fluorescence microscopy images showing the levels of Bodipy-cholesterol accumulated in control, U18666A+Probucol, and U18666A+Probucol+AEA treated primary astrocytes. F) Bodipy-cholesterol quantification of images as shown in E demonstrate that the cholesterol buildup triggered by the inhibition of NPC1 and ABCA1 in primary astrocytes is alleviated by either AEA, 2AG, or CBD addition even in the presence of the inhibitors (U18666A+Probucol). These data suggest that cannabinoids act through NPC1 and ABCA1-independent pathways. Data are represented as Mean ± SEM. ****P-value < 0.0001.

Based on the observation that endocannabinoid treatment can alleviate cholesterol buildup in aged astrocytes, we conducted further experiments to investigate whether endocannabinoids could enhance cholesterol transport to neurons in co-culture with aged astrocytes. To address this, we performed the same experiments outlined in Figure 4A but introduced a third condition in which AEA was added to the aged astrocyte-neuron co-cultures. The uptake of Bodipy-cholesterol by neuronal cells was quantified from fluorescence microscopy images, as depicted in Figure 5C. Our findings demonstrate that AEA treatment effectively improves the transport of cholesterol from aged astrocytes to neurons, as neurons co-cultured with aged astrocytes in the presence of AEA incorporated significantly higher levels of cholesterol (Figure 5D; Control = 1.0000 ± 0.0377, GAL = 0.5032 ± 0.0288, GAL + AEA = 0.7938 ± 0.0302, p < 0.0001).

Finally, to assess whether cannabinoids mobilize cholesterol through rescuing NPC1 or ABCA1- dependent pathways, AEA, 2-AG, and CBD were added to astrocyte cultures loaded with Bodipy- cholesterol in the presence of the NPC1 and ABCA1 inhibitors, U18666A and Probucol. Our results indicate that all three cannabinoids were able to alleviate lysosomal cholesterol accumulation in astrocytes treated with U18666A and Probucol (Figures 5E and 5F), suggesting that cannabinoids mobilize cholesterol through alternative pathways independent of NPC1 and ABCA1.

## Discussion

Adequate levels of membrane cholesterol are crucial for proper neuronal function. Previous studies have demonstrated that as hippocampal neurons age, there is an increase in the expression of Cholesterol-24-hydroxylase or CYP46, leading to cholesterol loss from the plasma membrane ^[5,13]^. This cholesterol loss results in overactivation of the PI3K-Akt pathway ^[9]^, which interferes with the lateral diffusion and endocytosis of postsynaptic AMPA receptors, preventing the induction of long-term depression (LTD), a cellular process involved in memory formation ^[23,24]^. Additionally, the constitutive reduction of synaptic cholesterol observed during normal aging affects NMDA receptor-mediated activation of CaMKII and leads to epigenetic repression of BDNF, a gene essential for synaptic plasticity and memory formation ^[12]^.

In a recent study, Guix et al.^[14]^ demonstrated that cholesterol redistribution in cortical neurons aged in vitro is altered due to enhanced Akt-mTOR activity, leading to NPC1 degradation. This study also reported increased levels of miR33 in these aged neurons, contributing to the downregulation of NPC1 and sequestration of cholesterol in lysosomes. These findings suggest that increased cholesterol hydroxylation via CYP46 in aging neurons, combined with impaired cholesterol redistribution due to NPC1 decay, leads to a significant imbalance of membrane cholesterol.

Since neurons depend on cholesterol delivered by astrocytes ^[11]^, we investigated whether cholesterol trafficking is affected in aged astrocytes, potentially interfering with the delivery of ApoE-cholesterol to hippocampal neurons. Using an accelerated aging model in rat primary hippocampal astrocytes, we found that cholesterol accumulates in lysosomal (LAMP2+) compartments in these aged cells. The lysosomal cholesterol accumulation in aged astrocytes was observed through three different methods: addition of the cholesterol analog Bodipy-cholesterol to the cultures and imaging its distribution, staining of endogenous cholesterol with filipin, and transfection of astrocyte cultures with the cholesterol probe D4-mCherry. We also observed cholesterol accumulation in primary human astrocytes aged in vitro. Furthermore, to validate the accelerated aging model, we assessed Bodipy-cholesterol distribution in primary astrocyte cultures from young (2 months old) and old (24 months old) mice, and similar levels of accumulated cholesterol were observed in old astrocytes.

Our results clearly demonstrate that downregulation of NPC1 and ABCA1 occurs in aging astrocytes, likely due to increased levels of miR33, a microRNA that reduces cellular cholesterol efflux under low cholesterol conditions. Our findings indicate that the combination of inefficient ApoE-cholesterol export due to ABCA1 downregulation and impaired cholesterol redistribution resulting from reduced NPC1 levels leads to cholesterol accumulation in the lysosomal compartment of aged astrocytes. The cooperative nature of these events was demonstrated by pharmacological inhibition of ABCA1 and NPC1 in primary astrocytes. Notably, ABCA1 downregulation appears to be necessary for cholesterol accumulation in aged astrocytes, as aged astrocytes treated with the ABCA1 agonist Evodiamine accumulate lower levels of cholesterol in lysosomes. These results are in line with the work of Yu et al. ^[25]^, showing that NPC1 astrocyte-specific deletion does not impact behavioral phenotypes, CNS histopathology, or synaptic function in mouse models.

Most of the existing evidence suggests that ROS accumulation plays an important role in brain aging and age-related cognitive impairments ^[26]^, mainly due to damage to lipids, proteins, and cellular DNA, disrupting cellular function ^[27]^. Here, we show that oxidative stress leads to cognitive decline in aging due to the generation of a Niemann-Pick phenotype in astrocytes. We demonstrated that miR33 expression is triggered by oxidative stress in primary astrocytes, and both NPC1 and ABCA1, which are target genes of miR33, are downregulated by peroxide treatment in primary astrocytes. Since oxidative stress accumulation is a hallmark of aging, these findings provide mechanistic insight to explain why aged astrocytes acquire a Niemann-Pick phenotype.

Supporting our results obtained in aging models in vitro, our qPCR results suggest increased miR33 levels and reduced amounts of NPC1 in the hippocampal tissue of 20-month-old mice compared to 2-month-old individuals. Although the relative contribution of astrocytes and neurons to these results should be assessed, the data confirm that aging triggers a Niemann-Pick phenotype in mouse hippocampal tissue.

In this work, we found that cholesterol transport from astrocytes to neurons is impaired by aging. We observed that hippocampal neurons uptake lower levels of cholesterol from aged astrocytes compared to controls. This effect was even more evident in old neurons with a higher dependence on astrocytic cholesterol, reinforcing the idea that aging affects cholesterol homeostasis in astrocytes and neurons with a cumulative effect. The impaired cholesterol transport from astrocytes to neurons could be explained by the decreased levels of the ABCA1 transporter observed in aging astrocytes since this transporter is required for the release of ApoE-cholesterol complexes. In this aging context, the scenario would be aggravated in astrocytes containing ε4 alleles of ApoE since it has been reported that the cholesterol transport by ApoE ε4 has decreased efficiency ^[28]^. The aging astrocyte-neuron co-cultures used in this work would provide a straightforward model to test this hypothesis.

Finally, taking into account previous results from Galles et al. ^[21]^, where it was shown that feeding *C. elegans* with AEA or 2-AG is able to inhibit dauer formation caused by impaired cholesterol trafficking in Niemann-Pick C1 mutants, we demonstrated that the lysosomal cholesterol accumulation in aged astrocytes can be alleviated by the treatment not only with the endocannabinoids AEA and 2-AG but also with the phytocannabinoid CBD. Additionally, we showed that AEA treatment is able to improve the neuronal uptake of cholesterol from aged astrocytes. Thus, considering that reduced neuronal cholesterol impacts synaptic receptor function and synaptic plasticity, the fact that cannabinoids are able to restore the cholesterol transport from aged astrocytes to neurons would have important implications in the aging field since it would contribute to explain observations showing improved cognition in old individuals treated with phytocannabinoids ^[29,30]^.

The mechanisms involved in the improved cholesterol recycling by cannabinoids have yet to be determined. Our results show that these mechanisms would not involve rescued NPC1 or ABCA1 function since cannabinoids exert their effect even in the presence of inhibitors of these two proteins, Probucol and U18666A. How cannabinoids bypass the effect of reduced NPC1 and ABCA1 to reverse cholesterol accumulation in aging astrocytes is not known, but some clues could be taken from a recent work by Hernandez-Cravero et al. ^[31]^. Through genetic analysis in combination with the rescue of larval arrest induced by sterol starvation, it was found that the insulin/IGF-1 signaling (IIS) pathway and UNC-31/CAPS, a calcium-activated regulator of neural dense-core vesicle release, are essential for 2-AG-mediated stimulation of cholesterol mobilization. Dense-core vesicles seem not to be very abundant in astrocytes, as almost only 2% of the vesicles have been described in these cells ^[32]^. On the contrary, secretory lysosomes have been identified in astrocytes, where they fuse with the plasma membrane in a Ca^2+^-regulated manner ^[33–36]^. The cannabinoid-stimulated cholesterol secretion by one of these two calcium-dependent pathways is a matter for future studies.

## Materials and Methods

### Hippocampal Rat astrocyte primary cultures

Rat primary astrocyte cultures were prepared from postnatal 1 (P1) Wistar rats. Hippocampi were dissected in cold Hanks 1x solution, cut into small pieces with scissors and homogenized by pipetting. Cells were recovered by centrifugation in 15 mL Falcon tubes and resuspended in DMEM 1x + 10% FBS + 1% Penicillin-Streptomycin + 1% GlutaMAX. Hippocampi from 5-6 pups were plated per 75 cm^2^ flask bottle. Once the culture reached confluence (approximately 10 to 14 days after plating) the flasks were shaken at 800 rpm for 1 hour to remove microglial cells. Astrocytes were recovered by trypsinization and plated at 25,000 cells/cm^2^ in glass coverslips or plastic plates.

### Adult mice astrocytes primary cultures

To obtain a cell culture of astrocytes from adult C57BL/6J mice the animals were killed using an approved method of euthanasia. Subsequently, the brains were carefully extracted, and the hemispheres were dissected using ice-cold 1x Hank’s solution. The tissue was finely fragmented into small pieces using sterile scissors and incubated with trypsin in a controlled incubator environment at 37°C with 5% CO_2_ for 15 minutes. Following this step, the tissue fragments were transferred into a 15ml Falcon tube, centrifuged, and the supernatant was discarded. The tissue was gently disaggregated by pipetting up and down using a medium consisting of DMEM/F12 supplemented with 10% FBS, 1% Penicillin-Streptomycin, and 1% GlutaMAX, and then plated onto culture dishes at a density of 50,000 cells per cm2.

### Neuronal rat primary cultures

Primary cultures of hippocampal neurons were prepared from embryonic day 18 (E18) Wistar rats as described in ^[37]^. Hippocampi were dissected and placed into ice-cold Hank’s solution with 7 mM HEPES and 0,45% Glucose. The tissue was then treated with 0,05% trypsin EDTA; (Invitrogen; Life Technologies Co.) and incubated at 37°C for 16 min and then treated with DNase (72 μg mL-1; Sigma-Aldrich) for 1 min at 37°C. Hippocampi were washed three times with Hank’s solution. Cells were dissociated in 5 mL of plating medium (Minimum Essential Medium (MEM) supplemented with 10% horse serum and 20% glucose) and cells were counted in a Neubauer Chamber. Cells were plated into pre-coated dishes with poly D-lysine (Sigma-Aldrich) (750.000 cells in a 10 cm dish and 270.000 in a 6 cm dish) and placed into a humidified incubator containing 95% air and 5% CO_2_. The plating medium was replaced with equilibrated neurobasal media supplemented with B27 and GlutaMAX (Gibco; Life Technologies Co.). Hippocampal neurons were kept in culture for 7, 14 or 21 days in vitro (DIV).

### Neuron-astrocyte co-cultures

Rat primary astrocyte and neuronal cultures were prepared according to the previous description. For co-cultures astrocytes were plated on plastic dishes, while neurons were plated on glass coverslips with wax dots ^[37]^. Subsequently, the astrocyte culture medium was replaced with neuronal culture medium, and the coverslips with neurons were flipped on top of the astrocytes. The wax dots serve as “feet” to avoid direct contact between neurons and the glial feeder layer.

### Human astrocyte primary cultures

Human astrocytes were obtained from biopsies. The tissues were placed on a sterile plate and covered with cold Hanks 1x solution. Magnifying glasses, forceps and scissors were used to remove blood vessels and meninges while preserving the clear part of the tissue. The clean tissue was transferred to a new plate with trypsin (enough to cover the tissue) and cut into small pieces with scissors. Tissue fragments were incubated with trypsin for 15 minutes in the incubator (37°C, 5% CO_2_) and transferred to a 15ml Falcon tube with 1 or 2 ml of Hanks 1x solution. After centrifugation Hanks was completely removed using a micropipette and the tissue was disaggregated in 1ml of DMEM / F12 10% FBS, 1% Penicillin-Streptomycin by gently pipetting up and down. The homogenate was transferred to a 50ml Falcon tube with a Cell Strainer on top. The whole homogenate was plated into a T25 flask with 8-10 ml of DMEM/F12, 10% FBS, and 1% Penicillin-Streptomycin and incubated until 80% confluence. Astrocytes were then trypsinized and plated at 25,000 cells/cm^2^.

### Aging in vitro of primary astrocytes

To age astrocytes *in vitro* a D-Galactose-induced model of aging was utilized. For this purpose, 25,000 cells per cm^2^ were seeded in DMEM 1x culture medium supplemented with 10% FBS, 1% Penicillin-Streptomycin, and 1% GlutaMAX. The culture was allowed to reach approximately 50% confluence. Subsequently, the culture medium was replaced with DMEM 1x without D-Glucose but containing 10g/L of D-Galactose, along with 10% SFB, 1% Penicillin-Streptomycin, and 1% GlutaMAX. After 5 days in this culture medium, senescence was verified through staining to detect Senescence Associated β-Galactosidase (SA-β-gal).

### SA-β-galactosidase staining

The SA-β-galactosidase staining was performed following the protocol described in ^[16]^. Briefly, astrocytes plated on 12 mm coverslips were washed twice with PBS and fixed with 3.7% formaldehyde in PBS for 3 minutes. The cultures were then washed again twice with PBS, followed by the addition of the X-gal staining solution, and incubated at 37°C (not in a CO_2_ incubator). The blue color precipitate develops after 2 hours of incubation and reaches its maximum between 12 and 16 hours. The stained cultures were observed under a bright-field microscope (Zeiss Axiovert 200, Hamamatsu digital camera C4742-98).

The staining solution contained 1mg/mL of X-gal (Life Technologies-BRL), a 40 mM sodium phosphate/citric acid buffer (pH 6.0), 5mM potassium ferricyanide, 5mM potassium ferrocyanide, 150 mM NaCl, and 2 mM MgCl_2_.

### MTT assay

For MTT assay astrocytes were plated at 50,000 cells/well in a 24-well plate. Following the designated experimental time points for Galactose treatment, the culture medium was removed and cells washed twice with 1x PBS. Next, 400 µL of MTT solution/well was added (0.5 mg/mL MTT in DMEM 1x supplemented with 10% FBS), and incubated for 4 hours at 37°C with 5% CO_2_. Once the incubation period is complete, the supernatant was carefully removed and formazan crystals were resuspended in 200 µL isopropanol. Formazan was quantified by transfer of 50 µL samples in triplicate to an ELISA plate and absorbance measured at 570 nm.

### Live-Dead assay

Cell viability of D-Galactose treated cultures was assessed by the LIVE/DEAD® Viability/Cytotoxicity Assay Kit (Molecular Probes). For this assay primary astrocytes were grown on glass coverslips as described above and viability was measured at 0, 2, 5 and 7 days after D-Galactose addition.

### Bodipy-cholesterol labeling

To label the astrocytes with Bodipy-cholesterol (Avanti polar Lipids inc.) for microscopy, the cells grown on glass coverslips were incubated overnight (O.N.) with 1µM BODIPY-Cholesterol diluted in culture medium. Following the incubation period, the culture medium was carefully removed, and the cells were gently washed twice with 1x PBS to remove any excess of Bodipy-cholesterol. The cells were then fixed with 4% PFS for 20 minutes. After fixation, the cells were rinsed three times with 1x PBS to remove any residual PFS. Subsequently the cells were either mounted using Mowiol for fluorescence microscopy or subjected to immunolabeling using specific antibodies for further analysis. Microscopy images of Bodipy-labeled cells were taken within two days after labeling to prevent Bodipy-cholesterol diffusion.

### Cholesterol detection by D4-mCherry probe

The plasmid encoding the cholesterol probe D4-mCherry was described by ^[17]^. Rat primary astrocytes were transfected in a 24-well multiwell plate at confluence higher than 80%. A total of 0.5 μg of the pDEST40-mCherry-D4 plasmid/well was used. Briefly, the 0.5 μg of plasmid was mixed with the 2 μL of Lipofectamine 2000, each dissolved in 50 μL of Opti-MEM medium, and incubated for 5 minutes at room temperature. 100 μL mixture was then added to each well and incubated at 37°C with 5% CO_2_ for 24 hours. Finally, the transfected cells were fixed and used for immunofluorescence studies.

### Immunofluorescence

For immunofluorescence cells were washed three times with PBS and fixed in 4% (w/v) paraformaldehyde (PFA) in PBS at room temperature for 20 min. Cells were then rinsed three times with PBS and permeabilized by 0.2% (v/v) Triton X-100 in PBS during 5 min. Next, cells were washed again three times with PBS and then treated to reduce nonspecific staining by incubation for 1 h in a blocking solution of 5% (v/v) horse serum (HS). Primary antibodies were incubated overnight at 4°C in solution of 1% (v/v) horse serum (HS). Then, cells were washed three times with PBS and incubated with secondary antibody for 1 h at room temperature, at a dilution of 1:500 antibody in solution of 1% (v/v) horse serum (HS). Nuclei were stained with DAPI for 5 min, then cells were washed twice with PBS, and coverslips mounted with Mowiol.

The primary antibodies used for immunofluorescence were the following: LAMP2 mouse monoclonal (DSHB Hybridoma Product H4B4) dil. 1/100.

anti-Rab5 mouse (sc-46692, Santa Cruz Biotechnology, Inc. CA, USA; RRID:AB_628191) dil. 1:100.

anti-Rab11 rabbit (cat 71-5300, Invitrogen; RRID:AB_87868) dil. 1:100.

β3 Tubulin rabbit (Abcam Cat# ab11314; RRID:AB_297918) dil. 1:500.

GFAP mouse monoclonal antibody (Millipore Cat # MAB3402, RRID:AB_94844) dil. 1:500.

For filipin staining, cells were incubated with filipin (Sigma) 125 μg/mL in 1X PBS for 2 hs at room temperature protected from light.

Secondary antibodies:

Alexa 546 anti-mouse (Thermo Fisher Scientific) dil 1/500,

Alexa 546 anti-rabbit (Thermo Fisher Scientific) dil 1/500,

Alexa 488 anti-mouse (Thermo Fisher Scientific) dil 1/500

Images were obtained on an LSM800 ZEISS confocal microscope and then processed with ImageJ for fluorescence quantification and co-localization analysis.

For Bodipy-cholesterol quantification, total fluorescence/cell area was quantified in at least 50 cells/group of at least three independent cultures. Data from each independent culture were normalized by the mean value of the control group and joined for statistical analysis.

Colocalization analysis was performed by the Pearson’s coefficient using JACoP plugin (ImageJ software) from images of at least 20 cells/experimental group from at least three independent cultures.

### Drug treatments

U18666A (Sigma) and Probucol (Sigma P9672, stock 1000x in ethanol) were added to the culture medium at 4 µg/mL and 10 µM respectively and incubated overnight together with Bodipy-cholesterol.

Rescue of cholesterol accumulation were performed by adding the drugs to cultures previously incubated overnight with Bodipy-cholesterol and further incubated for 6 h at the following concentrations: Evodiamine (Sigma, stock 1000x in DMSO) 10 µM, Anandamide (Noladin) 2 µM, 2-acylglycerol (Noladin) 2 µM and Cannabidiol (Epixann 5%, Caillon Hamonet) 5 µM. H_2_O_2_ was added to the culture medium at 250μM and incubated for 3 h.

### RT-PCR

Hippocampal tissue and primary cells were homogenized with Trizol Reagent (Ambion / RNA Life Technologies Co.) and the RNA was extracted with chloroform, precipitated with isopropanol and washed with 70% ethanol following the standard protocol. Total RNA was quantified by absorbance at 260 nm using a Nanodrop ND-100 (Thermo Fisher Scientific Inc.). Retrotranscription to first strand cDNA was performed from 1 µg of total RNA using the RevertAid H Minus First Strand cDNA Synthesis kit (Thermo Fisher Scientific Inc.). 20 ng of synthesized cDNA were loaded in the qPCR reactions using qPCR Master Mix (Fast start Sybr green Master; Roche) in QuantStudio 3 (Thermo Scientific). The primers purchased to Macrogen were used at 0.5 μM final concentration. The housekeeping gene Gapdh was used as endogenous controls. Primers used:

NPC1 fwd sequence: 5’-AAAGGCTGCAACGAGTCTGT-3’

NPC1 rev sequence: 5’-ACATGGCGTCCAAACCTAAG-3’

ABCA1 fwd sequence: 5’-GTGTGGACCCATACTCTCGCA-3’

ABCA1 rev sequence: 5’-TCAGGTAGTAACCCGTTCCCA-3’

GAPDH fwd sequence: 5’-CTCCCACTCTTCCACCTTCG-3’

GAPDH rev sequence: 5’-CATACCAGGAAATGAGCTTGACAA-3’

miRNA 33 and U6 levels were quantified using the miRCURY LNA miRNA SYBR Green kit from QIAGEN, following the manufacturer’s instructions. The procedure consisted of two steps: a universal first-strand cDNA synthesis and a specific qPCR amplification and detection with SYBR Green. In the first strand synthesis, all RNAs were simultaneously polyadenylated using a poly(A) polymerase and reverse transcribed using oligo-dT primers. The PCR amplification utilized Locked Nucleic Acids (LNA), RNA analogs known for their improved thermal stability when hybridized to a complementary strand. The amount of input material for the RT reaction was optimized, resulting in final amounts of 150 ng of initial RNA obtained through TRIZol isolation. Reactions were assembled in a final volume of 10 µl, consisting of 7 µl of sample, 1 µl of 10X RT enzyme mix, and 2 µl of 5X RT reaction buffer. Retrotranscription was performed at 42°C for one hour, followed by incubation at 95°C for 5 minutes to inactivate the enzymes, and immediately cooled to 4°C. Samples were stored at 4°C for immediate usage or at −20 °C for short-term storage. cDNA was diluted 1:60 in RNase-free water according to the manufacturer’s instructions. Each reaction was performed in a final volume of 10 µl, containing 3 µl of diluted sample, 1 µl of specific primer (miRNA 33 or U6), 0.5 µl of RNase-free water, 0.5 µl of ROX Reference Dye, and 5 µl of 2X miRCURY SYBR Green Master Mix. qPCR was performed using a QuantStudio 5 thermocycler (Applied Biosystems). The cycling conditions were as follows: 1 cycle of 2 minutes at 95 °C for enzyme heat activation, followed by 40 cycles of denaturation at 95°C for 10 seconds, and combined annealing/extension at 56°C for 60 seconds. Subsequently, a melting curve analysis was conducted between 60 and 95°C. Fluorescence data in the green (SYBR) and red (ROX) channels were collected during the annealing/extension step of each cycle. Relative quantification was performed using the ΔΔCT method, utilizing at least two technical replicates from at least three independent cultures, with U6 serving as the internal control.

### Western blot

Hippocampal tissue or primary cells were lysed on ice with a solution containing 1M Tris-HCl, 1% Nonidet P-40, 150 mM NaCl, 5 mM EDTA, 1 mM sodium orthovanadate, 1 mM dithiothreitol, pH 7.4, protease inhibitor cocktail (Roche) and phosphatase inhibitor cocktail 2 (Sigma-Aldrich). Protein concentration was determined by Pierce® BCA protein assay kit (Thermo Scientific). Protein samples were electrophoretically resolved within 12%Tris-HCl gels and afterwards transferred to nitrocellulose membranes. Membranes were blocked for 1 hour and incubated overnight at 4°C with the NPC1 primary antibody (rabbit polyclonal, Novus Biologicals, UK, NB 400-148).

Peroxidase-conjugated polyclonal goat anti-rabbit IgG H+L (Advansta R-05072-500) was used as a secondary antibody at 1:5000. HRP conjugated anti beta-actin antibody (HRP-60008 Proteintech) was used as loading control. Bands were visualized with the Amersham ImageQuant 800 and quantified using ImageJ software.

### Statistical analysis

Statistical analysis were performed using the GraphPad Prism software. Fluorescence quantification data were compared either by Nonparametric Mann-Whitney test or multiple experimental groups were compared by two-way Anova and Tukey’s multiple comparison test. Western blots, statistical analysis was performed using the nonparametric Mann-Whitney test. Statistical analysis of RT-PCR and Western blot data were compared by non-parametric One sample t test using hypothetical value 1 for the control condition. Data are represented as mean ± SEM. ****P-value < 0.0001, ***P-value < 0.001, **P-value < 0.01, *P-value < 0.05.

## Aknowledgenments

We thank Gonzalo Quassollo, Centro de Microscopía y Nanoscopía de Córdoba (CEMINCO) for his support in confocal microscopy; Laura Montroull and Andrea Pellegrini for their assistance in the cell culture facility. We thank Dr. Cecilia Conde, Dr. Mariana Bollo, Dr. Pablo Helguera (Instituto Ferreyra) and Dr. Carlos G. Dotti (CBMSO-CSIC) for their valuable contribution in the experimental and theoretical discussion.

