## Supplemental Figure 1 for "Impaired cholesterol transport from aged astrocytes to neurons can be rescued by cannabinoids"

### Supp. Figure 1

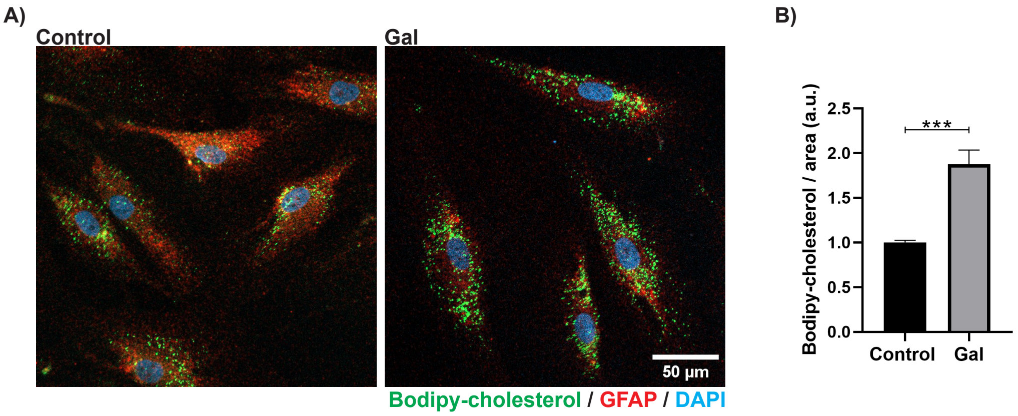


**Supp. Figure 1. Bodipy-cholesterol accumulates in human astrocytes aged in vitro.** A) Fluorescence microscopy images show that Bodipy-cholesterol accumulates in primary human hippocampal astrocytes aged by treatment with D-Galactose (GAL) compared to controls. Astrocytes were labeled with the astrocyte marker GFAP (red). B) Bodipy-cholesterol intensity quantification of images as shown in A (Control = 1.0000 ± 0.0233, Gal = 1.875 ± 0.160. Data are represented as Mean ± SEM. ***P-value < 0.001.
