## Supplemental Figure 2 for "Impaired cholesterol transport from aged astrocytes to neurons can be rescued by cannabinoids"

### Supp. Figure 2

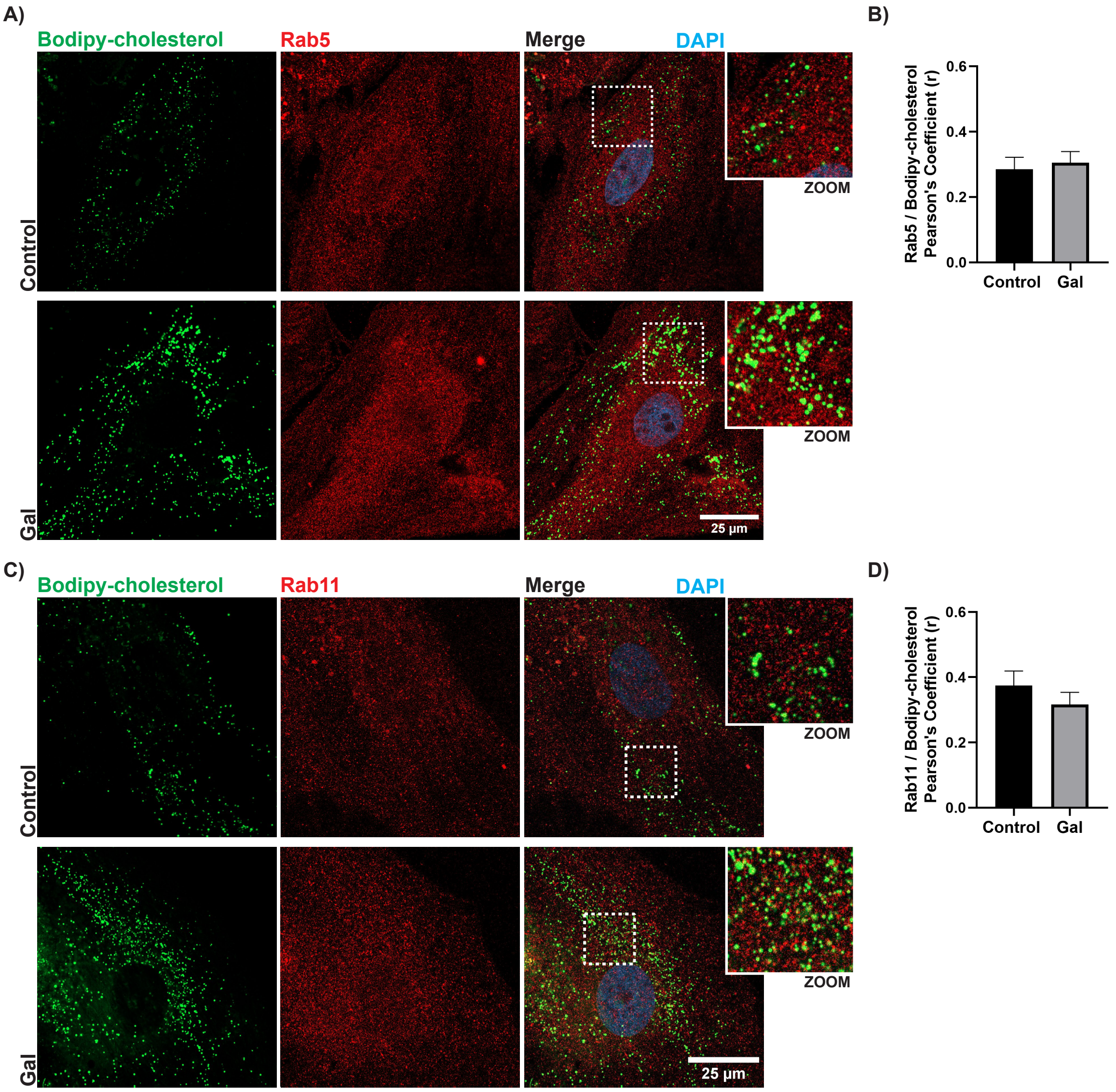


**Supp. Figure 2. Bodipy-cholesterol accumulated in aged astrocytes does not colocalize with Rab5 or Rab11 positive compartments.** A and C) Immunofluorescence confocal images of control or aged (GAL) rat hippocampal astrocytes labeled with Bodipy-cholesterol and the early (Rab5) and late (Rab11) endosomal markers. B and D) Analysis of images including those shown in A and C, demonstrates that the colocalization either between Bodipy-cholesterol and Rab5 or between Bodipy-cholesterol and Rab11 does not increase in D-Galactose treated astrocytes. Data are represented as Mean ± SEM.
