## Supplemental Figure 3 for "Impaired cholesterol transport from aged astrocytes to neurons can be rescued by cannabinoids"

**Supp. Figure 3**

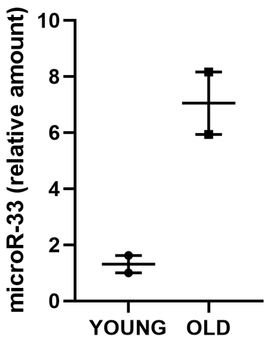


**Supp. Figure 3. miR33 increases during hippocampal aging in vivo.** Quantification of miR-33 levels by RT-qPCR in hippocampal tissue obtained from 2 young (2 monts-old) and 2 old (20 months-old) mice, indicating increased expression of miR-33 in hippocampal aging in vivo.
